# Genome-wide screen identifies new set of genes for improved heterologous laccase expression in *Saccharomyces cerevisiae*

**DOI:** 10.1101/2023.07.10.548373

**Authors:** Garrett Strawn, Ryan Wong, Barry Young, Michael Davey, Corey Nislow, Elizabeth Conibear, Christopher Loewen, Thibault Mayor

## Abstract

The yeast *Saccharomyces cerevisiae* is widely used as a host cell for recombinant protein production due to its fast growth, cost-effective culturing, and ability to secrete large and complex proteins. However, one major drawback is the relatively low yield of produced proteins compared to other host systems. To address this issue, we developed an overlay assay to screen the yeast knockout collection and identify mutants that enhance recombinant protein production, specifically focusing on the secretion of the *Trametes trogii* fungal laccase enzyme. Gene ontology analysis of these mutants revealed an enrichment of processes including vacuolar targeting, vesicle trafficking, proteolysis, and glycolipid metabolism. We confirmed that a significant portion of these mutants also showed increased activity of the secreted laccase when grown in liquid culture. Notably, we found that the combination of deletions of *OCA6*, a tyrosine phosphatase, along with *PMT*1 or *PMT2*, two ER membrane protein-O-mannosyltransferases involved in ER quality control, and *SKI3*, a component of the SKI complex responsible for mRNA degradation, further increased secreted laccase activity. Conversely, we also identified over 200 gene deletions that resulted in decreased secreted laccase activity, including many genes that encode for mitochondrial proteins and components of the ER-associated degradation pathway. Intriguingly, the deletion of the ER DNAJ co-chaperone *SCJ1* led to almost no secreted laccase activity. When we expressed *SCJ1* from a low-copy plasmid, laccase secretion was restored. However, overexpression of Scj1p had a detrimental effect, indicating that precise dosing of key chaperone proteins is crucial for optimal recombinant protein expression.

**Importance:** Our study showcases a newly developed high throughput screening technique to identify yeast mutant strains that exhibit an enhanced capacity for recombinant protein production. Using a genome-wide approach, we show that vesicle trafficking plays a crucial role in protein production, as the genes associated with this process are notably enriched in our screen. Furthermore, we demonstrate that a specific set of gene deletions, which were not previously recognized for their impact on recombinant laccase production, can be effectively manipulated in combination to increase the production of heterologous proteins. This study offers potential strategies for enhancing the overall yield of recombinant proteins and provides new avenues for further research in optimizing protein production systems.

## Introduction

The budding yeast, *Saccharomyces cerevisiae*, is a widely used host organism for the production of recombinant proteins, which include insulin, vaccines against HPV and hepatitis B, as well as various enzymes such as alpha-amylases and cellulases (1, 2). Even so, the historically low yields of protein from *S. cerevisiae* compared to other host organisms, often on the scale of milligrams of protein per liter of culture, potentially limits the value of such a system.

There are numerous bottlenecks which have the potential to severely hamper the capacity for recombinant protein production in *S. cerevisiae,* which include gene expression, correct folding of the protein within the ER, addition of post-translational modifications and trafficking for secretion (3). Overexpression of a recombinant protein may also trigger certain cellular stress responses such as the Unfolded Protein Response (UPR) and ER-associated degradation (ERAD) due to the accumulation of protein within the ER. Additionally, the overall metabolic burden can also limit the production of recombinant proteins. Engineering attempts to alleviate these bottlenecks have had varied results, with the success of an individual modification being largely dependent on the specific recombinant protein being expressed (4–6). Interestingly, recent work shows that modeling of the secretion pathway can guide the engineering of strains to increase recombinant expression (7). Nevertheless, no systematic screen has been done to confirm which cellular pathways are the main bottlenecks for heterologous protein production in yeast.

In this study we have utilized a fungal laccase enzyme, *ttLCC1*, isolated from *Trametes trogii* as our model recombinant protein (8). Laccases are multicopper oxidases with considerable biotechnological potential that can be found naturally in plants, insects, bacteria and fungi (9, 10). Their natural function varies depending on the organism, but can include lignification in plants and delignification in white-rot fungal species. Laccases from this group of fungi have received particular attention for their potential use in biotechnological applications due to their high redox potentials at the T1 copper site (8). Laccase enzymes show substrate promiscuity, capable of oxidizing a range of compounds including the common pollutant, Bisphenol-A, pesticides, and phenolic dyes (8, 9, 11–13). In addition, they are “green” enzymes in that they use readily available molecular oxygen and produce water as the only by-product. As a result of their many advantageous properties, laccases have been extensively studied for potential uses, as well as actual implementation in applications such as paper and pulp processing, synthetic chemistry, wastewater treatment, biofuel production from second generation feedstocks, and biofuel cells (14) and thus are of great interest for recombinant production.

To uncover additional potential engineering targets which can increase recombinant protein production in *S. cerevisiae*, we screened a library of 4,790 single gene, non-essential deletion mutants for effects on recombinant laccase expression and secretion using novel high throughput methodology based on solid media growth. We identified a first set of gene deletions that we further assessed in liquid cultures, resulting in several new gene deletions with significantly increased secreted laccase activity compared to a reference strain. This study showcases the use of novel high throughput methodology to identify novel engineering targets to increase recombinant production of laccases in *S. cerevisiae*.

## Results

### Screening of the laccase expressing single gene deletion mutants with the ABTS overlay assay and enrichment analysis of identified hits

To screen the yeast knockout (YKO) collection for effects on recombinant laccase production, a library of laccase-expressing single gene deletion mutants was generated using SGA methodology (15). The codon optimized *LCC1* gene from the fungi *Trametes trogii* under the control of the constitutively expressed and strong *GPD1* promoter with a N-terminal native secretion signal was integrated into the *TRP1* locus of the SGA query strain (JHY716). We verified that the activity of the secreted laccase could be readily assessed using a colorimetric assay, while no significant signal was noted from the original strain (Figure S1A, B). This laccase-producing strain served as the query strain for the SGA procedure and was mated to a collection of 4,790 unique gene deletions spanning the *S. cerevisiae* genome (Figure 1A). Three independent sporulations were performed in parallel before the multiple selection steps to generate the laccase-expressing deletion mutants and the library was decondensed (from 1536 colonies/plate) onto 48 plates at a density of 384 colonies per plate to facilitate high-throughput screening.

**Figure 1.**
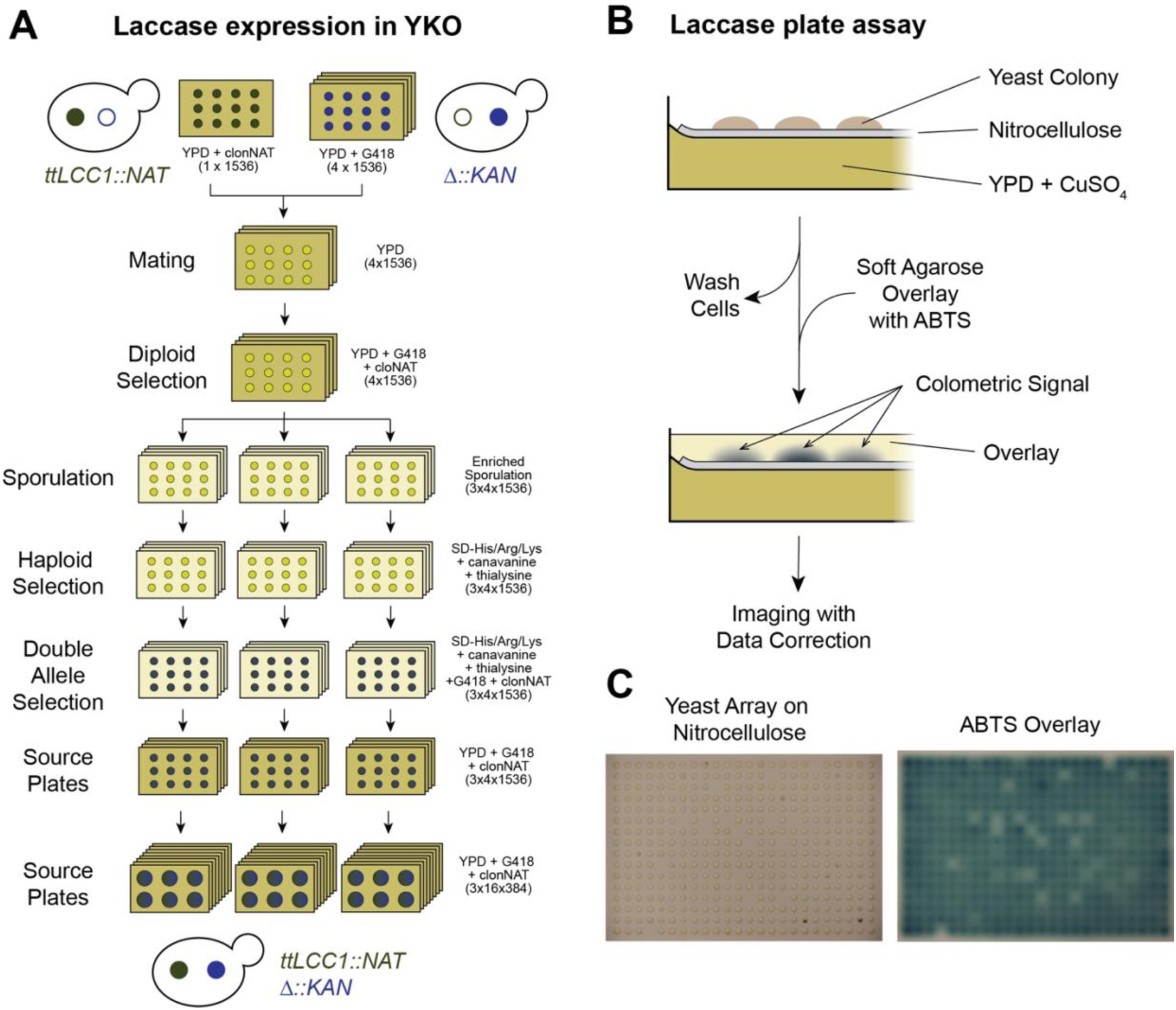
Screening of the YKO collection for laccase expression. **A.** Schematic of the establishment of a library with single gene deletion mutants expressing laccase using the SGA methodology. Approximately 5,000 strains were mated with the JHY716_*ttLCC1* query strain producing a total of 4,790 unique laccase-expressing single gene deletion mutant strains. **B.** Workflow of the ABTS overlay assay. **C.** Images of a representative array (plate 11, replicate 3) before washing (left) and an hour after addition of the overlay (right). Sites with differing amount of laccase activity can be identified by the intensities of the colorimetric signal.

To assess levels of recombinant laccase production and secretion from the generated library, a colorimetric ABTS (2,2’-azino-bis(3-ethylbenzothiazoline-6-sulfonic acid) overlay assay was developed (Figure 1B, C). High density yeast arrays were first pinned onto a nitrocellulose membrane overlaid on YPD media supplemented with copper (II) sulfate, which is necessary for proper folding and activity of the laccase enzyme (Figure 1B). To assess the levels of secreted laccase immobilized on the nitrocellulose membrane, cells were washed away from the membrane before the addition of a soft agarose overlay containing the ABTS substrate. One-hour post addition of the overlay, differing intensities were observed due to the varied activity levels of the secreted recombinant laccase from each YKO strain (Figure 1C). A custom image analysis pipeline incorporating densitometry was used to quantify the mean pixel intensity from each site on the assayed plate (see Methods). Importantly, normalizations to allow for inter-plate comparisons and corrections for the increased signal observed at sites near the peripheries of the plate were applied for the calculation of a modified Z score. Using this approach, we identified 66 “positive hits” with increase laccase activity and 208 “negative hits” with reduced activity among the 4,790 mutant strains that we assessed (Table S1).

A large portion of positive hits were mapped to the secretory pathway (Figure 2A) indicating their relevance during recombinant laccase production. In agreement with this observation, gene ontology (GO) analysis of the positive hits showed that several related processes; including Golgi retention, vacuole targeting, multivesicular body sorting pathway and vesicle transport are significantly enriched (Figure 2B). These results suggest that missorting of proteins could be a limiting factor in the production of recombinant laccase. Glycophosphatidylinositol (GPI) anchor biosynthesis was also an enriched GO term, as well as ATP export, autophagy and proteolysis. Intriguingly, deletion of the ER chaperone *LHS1* resulted in the greatest mean modified Z score (Figure S2A). Lhs1p has been shown to be a nucleotide exchange factor (NEF) of Kar2p, as well as being necessary for post-translational translocation into the ER lumen (16, 17). As a result, *lhs1*Δ mutants show a constitutive activation of the UPR (18). A constitutively active UPR could theoretically enhance the levels of recombinant laccase production through upregulation of other ER chaperones, expansion of ER size, and promotion of ER to Golgi transport. In contrast, deletion of *PET111* had the lowest mean modified Z score (Table S1). Pet111p is a translational activator for *COX2* mRNA which encodes for subunit II of Complex IV in the mitochondrial electron transport chain (19). Many genes involved in mitochondrial processes are observed in the list of negative hits suggesting the importance of functional mitochondria (Figure S2B). Correspondingly, there is an enrichment for numerous mitochondria-related GO terms among the negative hits (Figure 2C). Defective mitochondria could affect a variety of processes such as respiration and generation of ATP, maintenance of redox state, amino acid and lipid metabolism, and synthesis of other metabolites including heme. Perhaps the most obvious interpretation is that the ability of cells to perform respiration, which occurs after glucose has been depleted in the growth media, is limited or abolished in these cells (20). Interestingly, respiration after glucose depletion has been proposed to be a stage of growth where protein folding, and thus recombinant protein production, is optimized due to elevated NADPH levels from ethanol metabolism that can reduce oxidative stress produced during folding within the ER (21). Interestingly, the ERAD pathway was also identified in the GO analysis (Figure 2C). This includes *HRD1* and *UBC7* that encode, respectively, an E3 ligase and E2 enzyme involved in targeting misfolded proteins (22), whose deletion was detrimental to laccase production. The effect of modulating ERAD is likely specific to each heterologous protein, as it was previously shown that deletion of these genes can be used to engineer cell factories (23). Taken together, results from the ABTS overlay assay screen and subsequent enrichment analysis suggests that missorting during vesicle trafficking is a potential major limiting factor, and thus engineering target, for recombinant laccase production. It appears, from the negative hits, that functional mitochondria are beneficial for the production of recombinant laccase, possibly by promoting conditions that improve protein folding during respiration. Additionally, proteins involved in ERAD appear to be potentially necessary to sustain high levels of secreted laccase.

**Figure 2.**
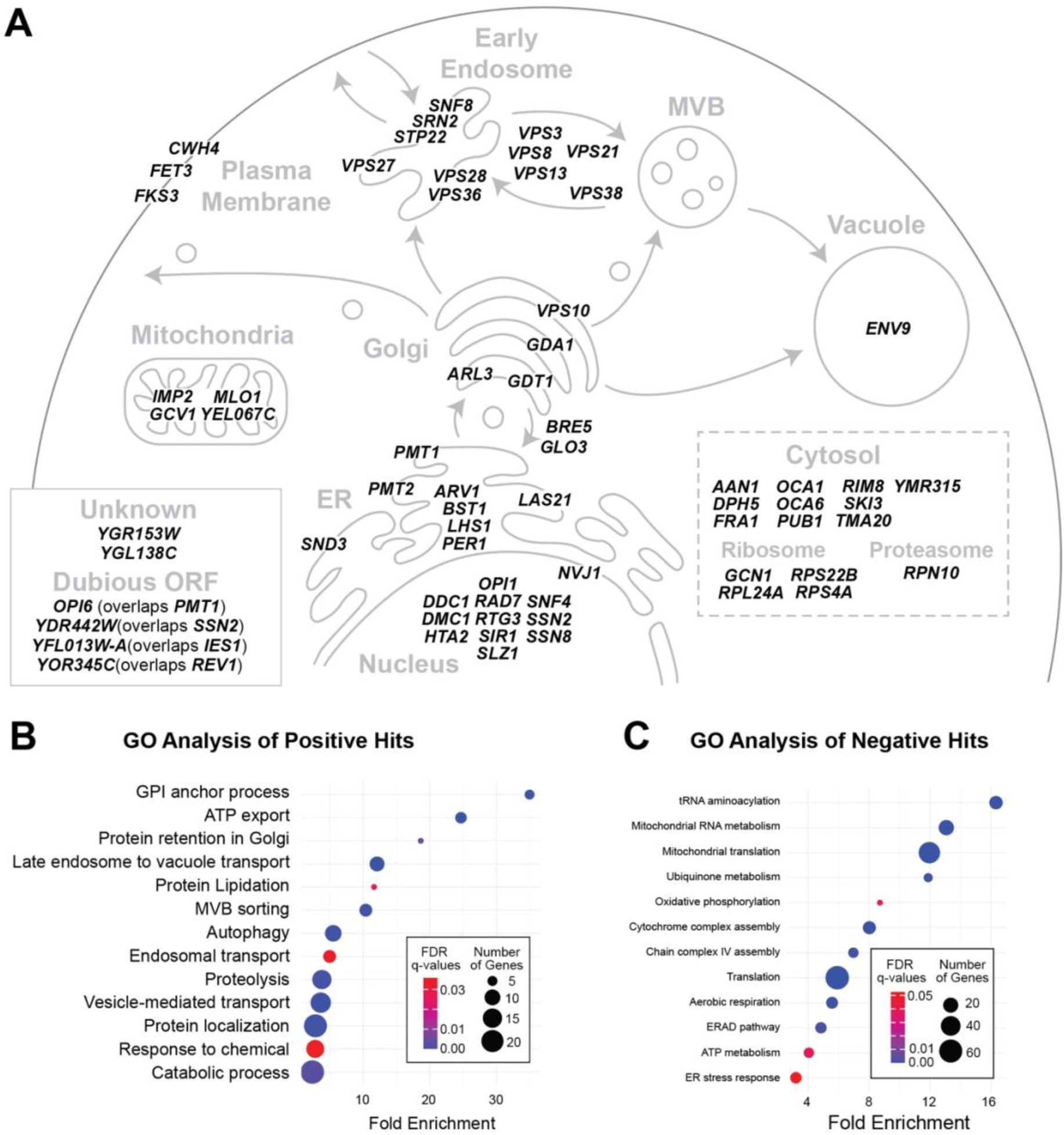
Enrichment of genes involved in vesicle trafficking and mitochondria. **A.** Representation of *S. cerevisiae* secretory pathway showing the cellular location of identified gene deletions that show an increase in recombinant laccase activity in the ABTS overlay screen. **C and D.** Dot plots of the main pathways enriched in the GO analysis of genes that lead to increase (C) or decrease (D) laccase activity in the YKO collection.

### Characterization of hits from ABTS overlay assay with liquid cultures

To further assess the increased laccase production observed in the ABTS overlay assay screen, the sixty-six identified positive hits were assayed in liquid culture, which is more similar to cultivation conditions when recombinant proteins are produced in bioreactors. Sampling of the cleared supernatant containing secreted laccase was performed at 96 hours post inoculation of the batch culture in 96 deep-well plates and quantification of secreted laccase activity was accomplished using a plate reader. A secreted laccase activity (µM oxidized ABTS / min) was calculated from the rate of change in absorbance using the Beer-Lambert Law. The OD_600_ of the microvolume cultures were measured in parallel and used to normalize the secreted laccase activity to account for the number of cells in each culture. The three biological replicates of the sixty-six positive hits were assayed alongside the reference query strain, JHY716_*ttLCC1*, to determine the change in normalized laccase activity from that of the parent strain. Out of the 66 positive hits, 50 were observed to have an increased normalized laccase activity in comparison to the reference strain, including 17 gene deletions with significantly increased normalized laccase activities (Figure 3A). Similar results were obtained when only the laccase activity was considered, regardless of the culture density (Figure S3A). All 17 gene deletions with significantly increased normalized activities had over a twofold increase in comparison to the reference strain with *ski3*Δ*, arv1*Δ, *pmt2*Δ strains each having fold increases of 5.3, 5 and 4.3, respectively (Figure S3B). Ski3p is a scaffold protein that is part of the cytosolic SKI complex that associates with the exosome to facilitate 3’-5’ mRNA degradation (24). Notably, we detected a significant increase of *LCC1* transcript levels in *ski3Δ* cells (Figure S3C). Arv1p is an ER membrane-localized flippase that is thought to be responsible for the transport of GPI intermediates into the ER lumen from the cytosol and loss of Arv1p disrupts organelle integrity and induces the UPR (25). Pmt2p is an O-mannosyltransferase that participates in ER protein quality control (26). Interestingly, deletion of several genes involved in late-stage vesicle trafficking such as *STP22, SRN2, VPS27,* and *VPS28* did not lead to an increase of laccase production in liquid culture while deletion of these genes displayed the greatest laccase activity increase in the solid media assay (Figure 3A, Figure S2A). Different cultivation conditions in the two assays may therefore differentially impact secretion of the recombinant protein.

**Figure 3.**
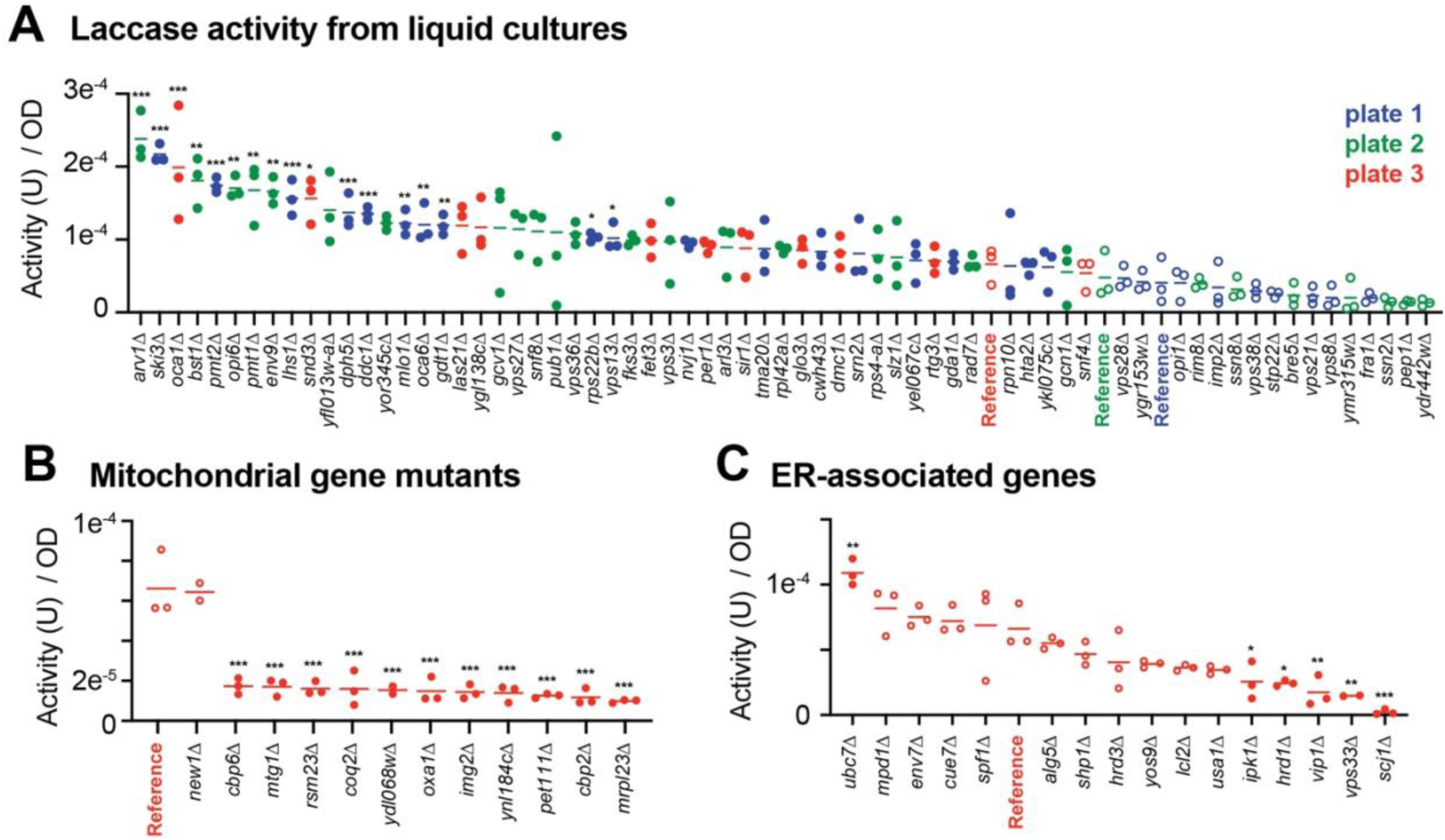
Change of laccase activity confirmed in liquid culture. **A.** Positive hits identified in the first overlay screen were assessed in liquid cultures. The strains were grown and assessed on three separate sets (plates 1-3) and ranked based on the average laccase activity (µmol/min) normalized to cell density. For each strain, three biological replicates were grown on the same plate and laccase activity was compared to the respective parental query strain (i.e., grown on the same plate) using ONE way ANOVA tests with Dunnett’s corrections (adjusted p values; < 0.05: *; <0.01: **: <0.005: ***). **B and C.** A subset of negative hits were assessed in liquid cultures. Three biological replicates for each strain were assessed and ranked based on the average laccase activity normalized to cell density as in A.

We next assayed a selected set of gene deletions from the 207 negative hits to validate their detrimental effect on recombinant laccase production. Deletions of select genes involved in mitochondrial processes were observed to have significant decreases in laccase activity, which also coincided with a reduced fitness of these mutants (Figure 3B, S3D). We then investigated negative hits with particular attention on proteins localized to the ER or involved in ER stress response pathways. In this case, only the *vip1Δ* strain displayed a significantly lower cell growth (Figure S3E) and most of the assessed mutant strains display a reduced activity, consistent with the overlay screen results (Figure 3C). Notably, deletion of the ERAD E3 ligase Hrd1p led to significantly reduced laccase activity. Similarly, deletion of several other genes associated with ERAD (*SHP1, HRD3, YOS9, USA1*) showed a decrease of laccase activity, albeit not in a significant manner. Deletion of two genes encoding for kinases involved in inositol processing, *IPK1* and *VIP1*, resulted in significantly decreased normalized activity. The same observation was made for deletion of HOPs and CORVET complexes subunit *VPS33*. Interestingly, deletion of *SCJ1* that encodes the ER HSP40 DNAJ co-chaperone almost completely diminished normalized laccase activity with no detrimental effect on cell fitness. This result suggests that the presence of Scj1p may be necessary for the proper production of the recombinant laccase. Taken together, these results indicate that most gene deletions that impact laccase activity in the overlay assay also affect recombinant laccase production in liquid culture.

### Rescue experiments to confirm hits

We performed a series of validation experiments by rescuing the deletion phenotypes with a plasmid-expressed form of the wild type gene. We first focused on *SCJ1* where the reintroduction of the gene with endogenous flanking sequences (including promoter and terminator) on a low-copy plasmid was sufficient to nearly restore the levels of the secreted laccase activity in comparison to the reference strain (Figure 4A). Interestingly, overexpression of *SCJ1* by using a high-copy 2µm plasmid only partially rescued the deletion phenotype (Figure 4B). Similarly, overexpression of the co-chaperone in the reference strain that contains a wild-type copy of *SCJ1* also resulted in a reduction of secreted laccase activity. This indicates that endogenous *SCJ1* likely has an optimal expression level for recombinant laccase production, and higher expression of this key gene was not able to increase the secreted laccase activity, but rather, limited it.

**Figure 4.**
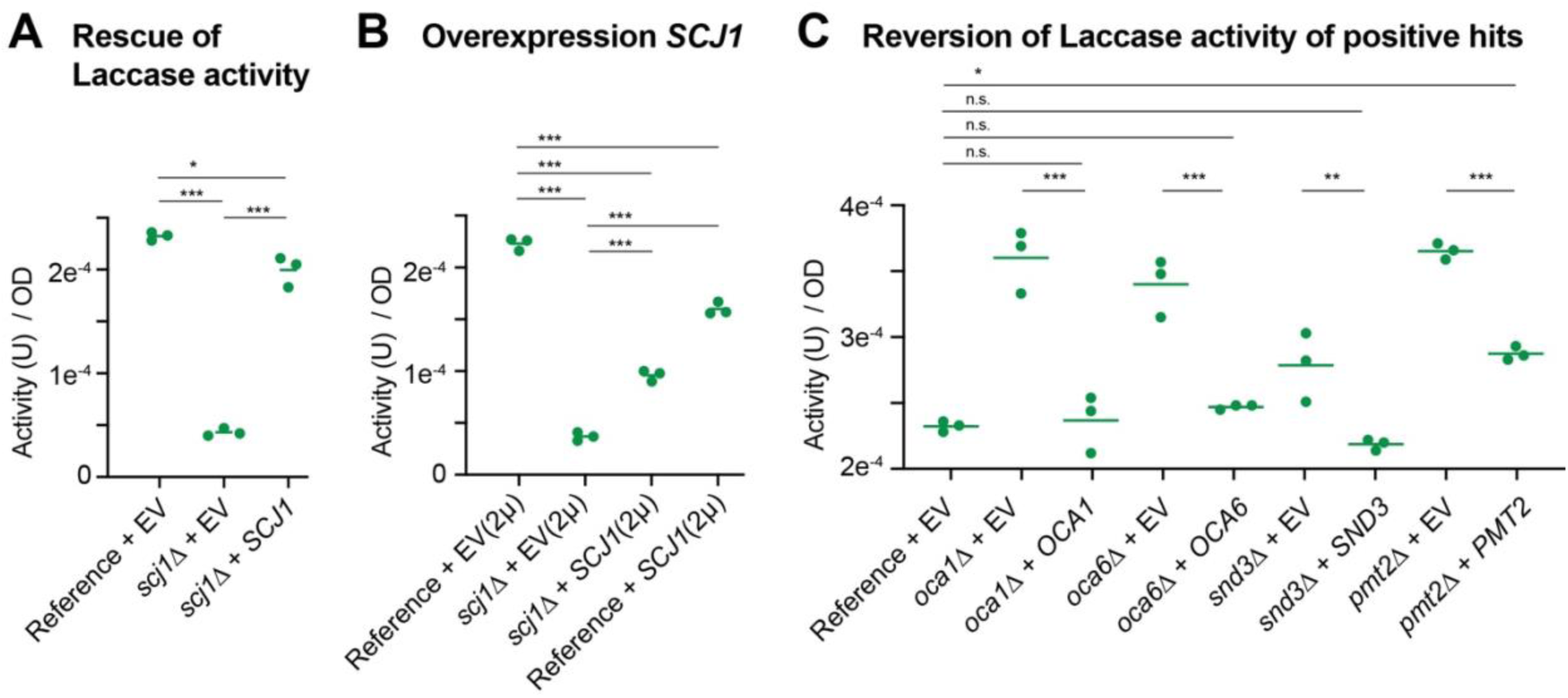
Rescue experiments in liquid culture. **A-C**. Normalized laccase activity of the indicated strains that contain either a CEN base plasmid (A and C) or a 2µ high copy plasmid (B) with the indicated genes subcloned with their 5’ and 3’ UTR regions or the empty vector (EV). ONE way ANOVA tests were performed with Tukey’s corrections (adjusted p values; < 0.05: *; <0.01: **: <0.005: ***).

We next subcloned ten different genes (*ARV1, BST1, ENV9, LHS1, OCA1, OCA6, PMT1, PMT2, SKI3, SND3*) into the low-copy plasmid and reintroduced them into the gene deletion strains. From these complementation experiments, expression of *OCA1*, *OCA6*, and *SND3* fully abrogated the gain of laccase activity in the mutant strains back to the reference strain levels (Figure 4C). In addition, expression of *PMT2* led to a significant decrease of laccase activity when compared to the *pmt2*Δ strain. The observed increase in secreted laccase activity could not be rescued for the remaining six ORFs (data not shown). Mutations already present in the deletion strains or accumulated during the library generation could be responsible for failure to rescue the high secreted laccase activity phenotype observed in the deletion strains. Alternatively, improper expression from the low-copy plasmid could result in a failure to reverse the observed phenotype.

### Double deletion screen

As several genes identified in our screen do not seemingly have overlapping functions, we sought to determine which double deletion mutants could lead to a higher laccase secretion. Using the JHY716_*ttLCC1* query strain, we deleted a subset of eight target genes by homologous recombination with the hygromycin drug resistance marker (hphMX). Using the SGA approach, we mated the newly established deletion strains with nine original deletion strains from the YKO selection then selected haploids cells with both deletions. We then assessed a subset of 50 different double mutants for laccase production in liquid culture that we normalized to cell density (Figure 5A). The average laccase activity was in general higher in comparison to controls where the hygromycin gene was integrated in the *HO* locus (first column) and several double mutants displayed over a two-fold laccase activity in comparison to the reference strain. This was the case for a cluster of strain combinations in which *OCA6*, *PMT1*, *PMT2* and *SKI3* were deleted (Figure 5A). In contrast, deletions of *ARV1* and *LHS1* often led to lower laccase activity in the double mutant strains. Because the mini-array was assessed on multiple plates and lacked some controls, we repeated this experiment with only a subset of the promising double mutants along with appropriate single deletion controls. Deletion of *OCA6* with *PMT1* or *PMT2* led to significantly higher laccase activity in several combinations in comparison to controls (Figure 5B). In contrast, double deletion of *PMT1* and *PMT2* did not further impact laccase activity when compared to single deletions. Deletion of *SKI3* also led to an increase of the average activity, albeit not always in a significant manner. These results show how the SGA technology can be used to screen many different combinations of mutations for strain optimization.

**Figure 5.**
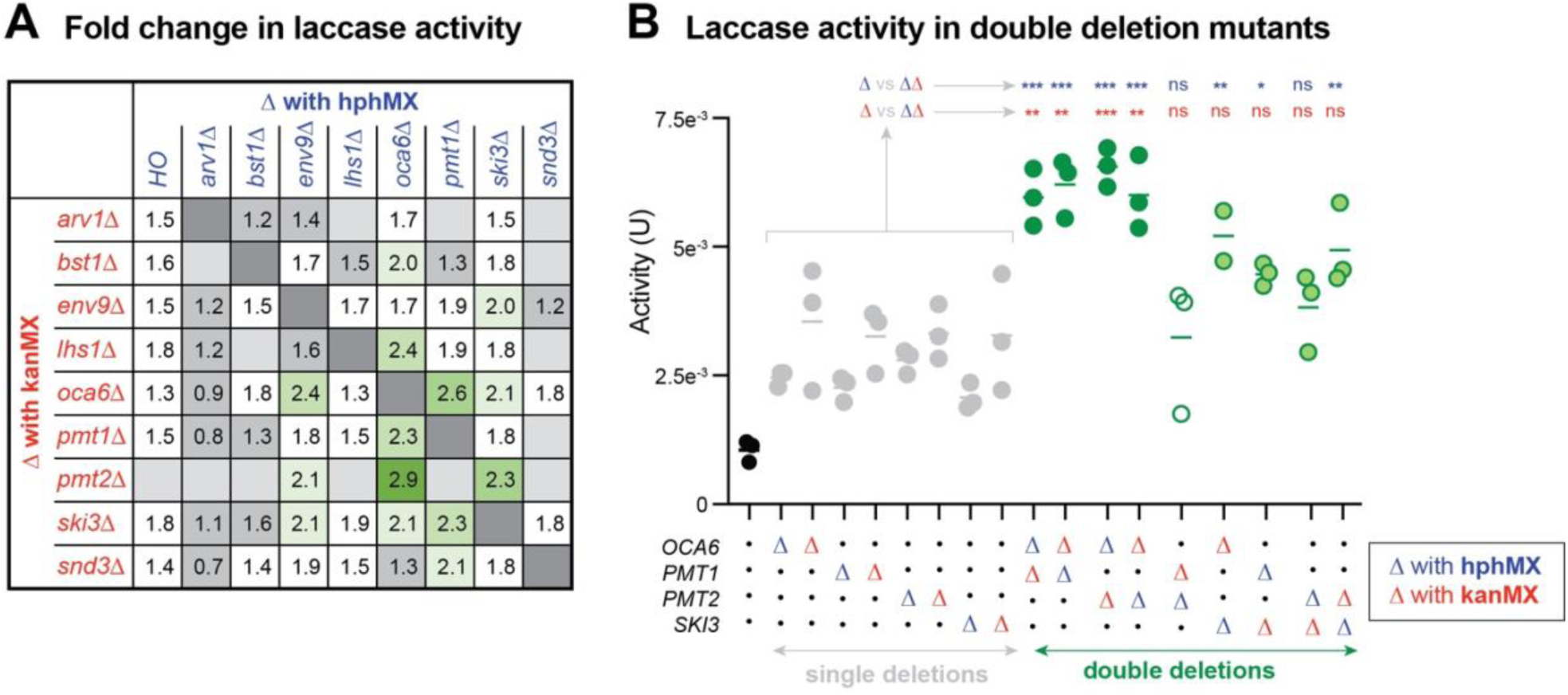
Double deletion mutants with increased laccase activity. **A.** Laccase activity of the indicated strains from a double deletion mini-array were assessed on three 96-well plates using the ABTS liquid culture assay after 4 days of growth. Three biological replicates were assessed for each double mutant with additional controls. Normalized laccase activity was averaged and compared to the activity levels from the reference strain. Gene deletions were either done by integrating the hygromycin (hph; blue) or kanamycin (kan; red) resistance modules. **B.** Laccase activity of the indicated strains grown on the same 96 well plate with three biological replicates. ONE way ANOVA tests with Tukey’s correction were performed and adjusted p values are shown for comparisons between double deletion mutants and corresponding single deletion mutant strains integrated with either the hph (blue) or kan (red) modules (< 0.05: *; <0.01: **: <0.005: ***).

## Discussion

In this study, we initially screened 4,790 unique gene deletions using the ABTS overlay assay and identified a set of 66 gene deletions that increased secreted laccase activity. Subsequently, we demonstrated that a significant proportion of these hits resulted in elevated activity of secreted laccase in liquid culture. We validated a subset of these mutant strains using a plasmid complementation approach. Lastly, we conducted a mini-array screen to identify double mutant strains with enhanced laccase activity.

Several of our identified hits in the genome-wide screen have been previously identified in other studies aiming to improve recombinant protein production or to study protein secretion in yeast. These include deletions of *ARV1*, *PER1*, *SNF8*, *VPS3*, *VPS27*, and *VPS28* in a solid media-based screen for increases in recombinant cellulase secretion (27). Additionally, *vps4*Δ, *vps8*Δ, *vps13*Δ, and *vps36*Δ strains have been shown to exhibit increased secretion of an insulin fusion protein (28). Deletion of the protein sorting receptor, *VPS10*, was identified in our screen and is also a common engineering strategy that has been employed previously in *S. cerevisiae* (29). Deletion of the negative regulator of lipid biosynthesis, *OPI1*, was also able to increase production of full-length antibodies four-fold (30). A number of the identified gene deletions have also been identified in screens that are not focused on recombinant protein production, but rather for defects in the secretory pathway including improper vacuolar sorting and ER homeostasis (31, 32). Such genes include the VPS genes identified in our screen (*VPS3, 8*, *13, 21, 27, 28, 36, 38*) and others (*ARL3, BRE5, BST1, GDA1, GLO3, LAS21, LHS1, PER1, PMT1, PMT2*) that have been identified in a screen for mutants that secrete CPY due to defects in vacuolar protein sorting (31). This is relevant as, under normal conditions, CPY is targeted to the vacuole for degradation suggesting that deletions of the above genes could also result in rerouting the recombinant laccase from vacuolar degradation to secretion. Additionally, *BRE5, BST1, LAS21, PMT1, PMT2, PER1* and *SND3* were identified in a screen for gene deletions that result in improper ER retention and secretion of the chaperone Kar2p (32). Along with the GO enrichment analysis, these results confirm that alteration of genes involved in intracellular protein transport can play a major role in heterologous protein expression in *Saccharomyces cerevisiae*.

We provided additional evidence to support the effect on laccase production of several genes that have not been previously implicated in recombinant protein production including the protein-O-mannosyltransferases *PMT1* and *PMT2*, a pair of phosphatases *OCA1* and *OCA6*, a subunit of the cytoplasmic SKI complex, *SKI3,* and *SND3* involved in ER targeting. These genes represent a set of engineering targets that could be applied in future studies. Most promising are the combined deletions of *OCA6* and *PMT1* or *PMT2* that led to higher increase of activity of the secreted laccase, and potentially the deletion of *SKI3*.

Oca1p and Oca6p are phosphatases that localize to the cytoplasm. Oca1p is known to associate with other oxidant induced cell cycle arrest (OCA) proteins in a complex, while Oca6p does not appear to be a part of the complex (33). Both their biological roles are relatively unknown; however, recent investigations have uncovered links between inositol metabolism as well as translation initiation when OCA genes are deleted.

Possible explanations of how deletion of *PMT2* results in an increase in recombinant laccase production include the observation that deletion of *PMT2* results in the failure of functional unfolded protein O-mannosylation (UPOM) (34, 35). A non-functional UPOM would presumably allow for increased folding cycles of the recombinant laccase by the Kar2p chaperone. Additionally, abolition of O-mannosylation through deletion of *PMT2* has been shown to result in decreased cell wall integrity as the majority of cell wall proteins are heavily mannosylated (36). Thus, if secreted laccase was unable to diffuse past the cell wall, decreased cell wall integrity could increase levels of secreted laccase in the culture supernatant.

Ski3p serves as the scaffolding subunit of the SKI complex, a cytoplasmic complex that is involved in the 3’-5’ degradation of normal mRNAs, non-sense mediated decay, and non-stop mediated decay (37–39). Consistent with a possible role of the SKI complex in regulating expression of the recombinant protein, levels of *LCC1* mRNA were significantly higher in *ski3Δ* cells. Interestingly, deletion of other SKI complex genes such as *SKI7* and *SKI8* led to increased levels of secreted laccase by the ABTS overlay screen with *ski8*Δ having a mean modified Z-score of greater than 2.5 but were not classified as hits due to variability between replicates. Deletion of the last SKI complex gene, *SKI2*, which is an RNA helicase, only resulted in a slight increase of secreted laccase activity, while deletion of the SKI2-like helicase, *SLH1*, resulted in secreted laccase activity comparable to that of deletion of *SKI7* and *SKI8*, possibly suggesting a similar effect to deletion of *SKI3, SKI7* and *SKI*8.

*SND3* encodes for a protein that is involved in SRP-independent post-translational translocation (40). Therefore, the increased laccase activity in *snd3*Δ is potentially caused by an overall decrease in the number of proteins within the ER and post ER vesicles. Alternatively, it has been recently shown that Snd3p plays an essential role in mediating the expansion of perinuclear ER-vacuole junctions (NVJs) during glucose starvation (40). Lipid droplet biogenesis occurs at these NVJs upon glucose starvation (41). Abolition of NVJs by the deletion of *SND3* could therefore prevent an increase of lipid droplet formation and limit the loss of phospholipids from the ER membrane during glucose starvation, thus potentially allowing for increased vesicle formation at ER exit sites and secretion of the recombinant laccase. Interestingly, deletions of *VSP13* and *NVJ1*, which encode for a lipid transporters at membrane contact sites and a tether protein that also mediates NVJs, respectively (42, 43), were also identified in our screen. This suggests that abolishing NVJs is an effective strategy to increase recombinant laccase production possibly through modulation of lipid droplet formation. In agreement with this possibility, deletion of *ENV9*, which encodes for an oxidoreductase that is involved in lipid droplet morphology (44), also led to higher laccase activity. Deletion of *ENV9* results in decreased lipid droplet size (45) which, again, could allow for a higher proportion of the ER membrane to be utilized for vesicle formation. Additionally, the *env9*Δ null mutant shows defective vacuole morphology, which could also possibly explain the increase in enzyme laccase activity if a portion of the recombinant laccase is normally directed towards the vacuole for degradation.

In addition to positive hits, we identified 207 negative hits that showed decreased laccase activity. The majority of the negative hits were genes involved in mitochondrial processes, which presumably resulted in decreased cell fitness, and thus was detrimental to recombinant protein production. Several ER-localized genes involved in ERAD (e.g., HRD1, *SHP1, HRD3, YOS9,* USA1), N-linked glycosylation (*ALG5, ALG8, DFG10*) translocation (*GET2*) and protein folding (*MPD1* and *SCJ1*) were identified, suggesting the importance of these processes during recombinant laccase production (46–49). Interestingly, while we could rescue the loss of laccase activity in *scj1*Δ cells by expressing *SCJ1* on a low copy plasmid, higher expression of this gene was detrimental. Indeed, it has also been previously observed that overexpression of *SCJ1* resulted in a decrease in the production of recombinant human albumin in log phase *S. cerevisiae* cultures (50). Therefore, careful dosage experiments should be performed when engineering host strains with supplementary copies of chaperone proteins.

### Limitations of our study

One limitation of the overlay screen is that yeast cells are most commonly grown in fed batch fermenter tanks rather than on solid media. Indeed, a few positive and negative hits show opposite results when re-assessed in liquid media (e.g., *VPS8* and *UBC7*). However, using the presented assay, we were able to screen approximately 15,000 total strains for their level of recombinant laccase production in just under a week once the library was generated (∼5000 strains in 3 biological replicates).

## Methods

### Plasmid and Yeast Strain Construction

All plasmids, yeast strains and oligos used or generated in this study are listed in Table S2. To construct a query strain for the SGA procedure, the integration vector YIp-*TRP1*-natMX, a gift from Dr. Hampton, was digested with BamHI and SacI to insert the codon optimized *ttLCC1* gene flanked by the constitutive *GPD1* promoter and the *CYC1* terminator (CYC1t) from pRS314-*ttLCC1*-natMX (BPM1768) to generate YIp-*TRP1*-*ttLCC1*-natMX (BPM1843) (51). The original source of codon optimized *ttLcc1* with a N-terminal native secretion signal was a gift from Dr. Sychrová (8). The YIp-*TRP1*-*ttLCC1*-natMX plasmid was linearized with the Bsu36I restriction enzyme in the *TRP1* gene and integrated into the JHY716 strain (*MATα, can1*Δ*::STE2pr-Sp_his5, lyp1*Δ, *his3*Δ*1, leu2*Δ*0, ura3*Δ*0, met15*Δ*0, cat5(I91M), SAL1, mip1(A661T), HAP1, mkt1(D30G), rme1(ins-308A), tao3(E1493Q)*) at the *TRP1* gene via a high efficiency LiOAc transformation protocol (52). Successful integration was confirmed *via* PCR using a primer pair specific to both the genomic DNA 5’ of the integration site and integrated DNA (Table S2) and the generated query strain was named JHY716_*ttLCC1* (YTM2204).

To create independent gene knockouts of selected hits, an hphMX drug resistance cassette encoding for resistance against the antibiotic hygromycin B, was amplified from pAG32 (53). Primers used included 40 bp of homology to the immediate 5’ and 3’ untranslated flanking sequences, including the start and stop codons, of the targeted open reading frame (ORF) (Table S2). The same high-efficiency LiOAc transformation protocol as above was used for integration of PCR amplified DNA (52). Integration was confirmed *via* a colony PCR protocol using primers specific to the surrounding genomic sequence and the integrated hphMX DNA (Table S2).

For complementation experiments, DNA was PCR amplified from JHY716 genomic DNA. Primers were designed to specifically amplify the ORFs and flanking sequences, that included annotated transcription start sites, TATA like elements, 5’ UTRs, and 3’ UTRs, from genomic DNA (Table S2). Homologous sequences to modified pRS416 (BPM1745) and modified pRS426 (BPM1756), both containing hphMX instead of *URA3*, were used for Gibson Assembly following manufacturer’s instructions.

### Laccase Expressing Single Gene Deletion Library Generation and ABTS Overlay Assay Screen

To construct a genome wide library of laccase expressing single gene deletion mutants, the synthetic genetic array (SGA) methodology was utilized (54). To start the SGA procedure, a 30 mL culture of the query strain, JHY716_*ttLCC1* (YTM2204), was grown in Yeast Peptone Dextrose (YPD) (2 % w/v) media at 30 °C with shaking overnight. The next day, a Singer Rotor HDA robot was used to array the query strain from the liquid culture onto a YPD + clonNAT (100 µg/mL) Singer plate in 1536 colonies per plate (cpp) density. Simultaneously, a recently pinned Deletion Mutant Array (DMA) collection also known as the YKO collection was condensed from 384 cpp to 1536 cpp on four YPD + G418 (200 µg/mL) Singer plates. Plates were incubated at 30 °C overnight. Next afternoon, the laccase expressing query strain was mated with each DMA plate by first pinning the query strain onto four different YPD Singer plates. Cells from the condensed DMA were then pinned on top of the query strain cells. Cells were allowed to mate for ∼18 hours before diploid selection by pinning cells onto YPD + G418 + clonNAT Singer plates. Cells were grown for ∼28 hours at 30 °C. Cells from each diploid selection plate were transferred onto 3 independent sporulation plates (1 % (w/v) Potassium Acetate, 0.1 % (w/v) Yeast Extract, 0.5 g/L dextrose, 0.1 g/L amino acid supplement powder (0.5 g histidine, 2.5 g leucine, 0.5 g lysine, 0.5 g uracil), 2 % (w/v) agar, 50 µg/mL G418) creating three biological replicates of the library (12 plates at 1536 cpp). Cells were grown on sporulation media for 5 days at 22 °C. Cells from the sporulation plates were transferred onto synthetic defined (SD; without ammonium sulfate and with 1 g/L MSG) – His/Arg/Lys + canavanine (100 µg/mL) + thialysine (100 µg/mL) Singer plates and grown for 2 days at 30 °C. Double mutants were then selected for by transferring cells onto SD – His/Arg/Lys + canavanine + thialysine + G418 + clonNAT plates. Plates were left at room temperature for ∼ 3 days. Cells were then transferred onto YPD + G418 (200 µg/mL) + clonNAT (100 µg/mL) plates at a density for 1536 cpp density for storage. Decondensing of 1536 cpp arrays to a density of 384 cpp on YPD + G418 (200 µg/mL) + clonNAT (100 µg/mL) plates was done to facilitate screening of the generated library using the ABTS overlay assay.

A similar approach was used to generate double deletion mutants on a mini-array. Strains were first spotted on two 384 cpp density YPD plates: one in which each column contains a different hph-integrated strain, and the other with each row containing a different kan-integrated deletion strain. Diploid selection was done on YPD + G418 (200 µg/mL) + clonNAT (100 µg/mL) + hygromycin B (200 µg/mL), diploid cells were spotted on three separate plates for sporulation, and following haploid selection, the final selection was done on SD – His/Arg/Lys + canavanine + thialysine + G418 + clonNAT + hygromycin B plates. Colonies were then transferred on YPD + G418 + clonNAT + hygromycin B plates before transferring cells to liquid cultures for the ABTS liquid assay. For each biological replicate, cells were derived from a different sporulation plate.

An ABTS overlay assay was used to screen the generated library. Arrays from SGA library preparation at a density of 384 cpp were pinned onto a nitrocellulose membrane (0.45 µm pore size) overlaid on YPD + G418 (200 µg/mL) + clonNAT (100 µg/mL) + CuSO_4_ (0.6 mM) media in Singer PlusPlates using a Singer Rotor HDA robot. Equal amounts of media, 40 mL, were deposited into plates to ensure equal focal length during imaging of the assay. Plates were incubated at 30 °C for 48 hours before colonies were washed from nitrocellulose membrane with a stream of phosphate buffered saline (PBS) solution. 40 mL of a heated solution (55 °C) of 0.5 % (w/v) agarose, 50 mM Britton and Robinson Buffer (0.1 M each of Boric, Phosphoric and Acetic Acid, brought to pH 4.0 with NaOH), and 0.5 mM ABTS was then administered onto the plate. The plate was incubated at room temperature for one hour to allow the colorimetric reaction to develop before images were taken with a digital camera (Canon Rebel EOS T3i, Manual Settings: f 2.8, 100 ISO, and 1/80 s exposure). An imaging setup built into the BM3-BC robot (S&P robotics) was utilized for consistent background illumination. Before images of the colorimetric reaction were captured, a blank plate overlaid with a nitrocellulose membrane was used to focus the camera. Images were stored as lossless RAW files (.CR2 file format) in addition to JPEG lossy file formats for visual inspection.

### Image Analysis and Quantification of Colorimetric Signal

A custom CellProfiler pipeline was used for image analysis and quantification of pixel intensity using densitometry from the colorimetric reaction. First, RAW images were cropped to defined dimensions, converted to grayscale, and then the pixel intensity was inverted for quantification purposes. A pre-prepared background illumination function was prepared by imaging six individual “blank” nitrocellulose plates overlaid with an agarose overlay. The images from these six blank plates were analyzed using CellProfiler to generate a background illumination function. Specifically, the minimum pixel intensity for each segmented section of the image (5 x 5-pixel dimensions) was found. A median smoothing filter was applied which removes bright or dim features that are imaging artifacts. The resulting background illumination function was subtracted from pixel intensities of images to subtract the background illumination from the nitrocellulose from the image. To quantify pixel intensity from the ABTS overlay assay, a 24 x 16 grid (384 total segmented blocks corresponding to the number of cpp) was overlaid onto the image in order to segment each site on the array for quantification. Within each segmented block a 101-pixel diameter circle, centered in each box on the grid, was used to define the boundary for quantification. The mean pixel intensity of pixels within the segmented circles was calculated for each site and exported to a spreadsheet for further data processing. We used a custom-built R script to correct for colony position due to the increase in signal intensity seen in the outer edges of the array. Importantly, before normalization and correction was applied, images of arrays before washing away of colonies were manually inspected to determine if growth occurred at each individual site, with a value of 1 representing “normal growth”, while a value of 0 represented “no growth” or “abnormal growth”. This manual inspection was necessary to prevent calculated plate medians to be influenced by a large number of empty sites with little to no signal. The mean pixel intensity for each site on an individual plate was then first normalized to the median mean pixel intensity of the plate, excluding sites with “no growth” or “abnormal growth”. To correct for increased signal of edge sites, a “zone correction” was then applied. This correction method consists of dividing the array into zones, defined as a series of concentric rectangles starting from the periphery of the plate and moving toward the center. If an individual zone had a median of the normalized mean pixel intensity values above the median of the mean pixel intensity of the entire plate (1.0 by definition), all normalized mean pixel intensity values within that zone will be divided by the median value of the zone. This is similar to “row” and “column” corrections used during the quantification of colony size in high density yeast arrays, however it is customized to the unique pattern of increased signal seen in our assay, resembling concentric rectangles (55). After normalization and zone correction of the mean pixel intensity for each site, modified Z-scores were calculated for each site in an individual plate. First median absolute deviations (MAD) were calculated using the formula *MAD* = 1.4826 ∗ *Median*(|*x* − *median*|). Modified Z-scores were then calculated by subtracting the median normalized and zone corrected mean pixel intensity from individual pixel intensity values and dividing by the calculated MAD. A threshold of modified Z scores greater or equal to 2.5 for 2 or more replicates was used to identify hits from the screen.

### Gene Ontology enrichment analysis and other statistical analyses

GO process enrichment analysis was performed using the online tool, GOrilla (56). The set of 66 positive hits and 207 negative hits were independently analyzed against a background list of the 4,790 gene deletions in the generated library using the Benjamini and Hochberg correction. All the statistical tests were performed in PRISM. Unless specified, ONE way ANOVA analyses were done with Dunnet’s correction for multiple comparisons to the same reference strain, and with Tukey’s correction for multiple comparisons between all the different strains.

### ABTS Liquid Assays

Selected hits from the previous screen were transferred from YPD + G418 + clonNAT source plates onto a new YPD plate and grown for 2 days at 30 °C. Cells were then inoculated into 150 µL of expression media (YPD, 20 µg/mL adenine, 50 mM potassium phosphate (dibasic) (pH 6),

0.5 mM copper (II) sulfate) in a 96 well round bottom plate and grown in a microplate shaker incubator at 30 °C with shaking at 900 rpm overnight. The next morning, culture density was measured with a plate reader (BMG, Clariostar +). Overnight cultures were then used to inoculate 500 µL of expression media at a starting OD_600_ of 0.2 in a 96 well 2 mL deep well plate. Deep well plates were sealed with a breathable cover and grown for 4 days (96 hours) at 30 °C with shaking at 900 rpm. At the end of 4 days, the OD_600_ of the cultures were measured again in order to normalize secreted laccase activity to the number of cells. The deep well plate was then spun down in a swinging bucket centrifuge at 3,200 rcf for 5 minutes to separate secreted laccase in the supernatant from the cells. 20 µL of supernatant containing the secreted laccase was then added to 80 µL of Britton and Robinson buffer (100 mM, pH 4) in a flat bottom 96 well plate. Immediately prior to quantification, 100 µL of 2 mM ABTS in 100 mM Britton and Robinson buffer (pH 4) was added to wells thus starting the colorimetric reaction. Secreted laccase activity was monitored over the course of two hours using UV-Visual spectrophotometry at 420 nm, the absorbance maximum of the oxidized ABTS product, with readings taking place every minute. Absorbance was plotted against time in order to determine the range where a linear rate of change is observed. Linear regressions were fitted to data with data points eliminated until a correlation coefficient of at least 0.999 was obtained. Using the Beer-Lambert law, absorbance was used to calculate the concentration of oxidized ABTS in µmols. A laccase activity value (µmols oxidized ABTS / min) was then calculated. The activity of value was normalized to the cultures OD_600_ (Activity / OD_600_) to control for differences in number of cells and were averaged across replicates for each strain.

To validate identified gene deletions, the replicate colony with the median normalized activity value was transformed with a plasmid containing rescue DNA specific to that gene. Simultaneously, an empty plasmid control, pRW113 (BPM1745), was transformed into the same gene deletion strain. Transformants were selected for on YPD + hygB (200 µg/mL) plates. Transformants were pooled together and spotted onto another YPD + hygB (200 µg/mL) plate as a source for future inoculations. When possible, the number of transformants pooled together was thirty. Assessment of secreted laccase activity was done as described above using expression media + hygB (200 µg/mL).

### RT qPCR

Total RNA was extracted using the RiboPure Yeast RNA Prep Kit (Thermo Fisher Scientific AM1926). RNA Integrity Number (RIN) and concentration was determined using the Bioanalyzer 2100 (Agilent G2939BA) and RNA 6000 Nano chip (Agilent 5067-1511). *ttLCC1* and *UBC6* mRNA levels were determined using the Power SYBR® Green RNA-to-CT™ 1-Step Kit (Thermo Fisher Scientific 4389986) using 70 ng of RNA and 10 µM primers (Table S2). Three 20 µL replicates were pipetted into a 384-well PCR plate, then sealed with an optically clear seal. RT-qPCR was run in a ViiA 7 Real-Time PCR System (Thermo Fisher Scientific 4453545) with cycle settings following manufacturer’s protocols for the RT-qPCR kit. Data was visualized and exported to Excel using QuantStudio Real-Time PCR Software (Thermo Fisher Scientific v1.6.1). Relative *ttLCC1* mRNA levels were calculated using the ΔΔCt method (57).

### Availability of data and materials

All data generated or analysed during this study are included in this published article and its supplementary information files. Additional datasets generated during the current study are available from the corresponding author on reasonable request. Plasmids and yeast strains listed in the supplemental material with a BPM or YTM denomination are available upon request.

### Authors’ Contribution

GS designed, executed and analyzed the data of most experiments. RW generated some plasmids, developed the liquid ABTS assay, designed and performed the RT-qPCR experiment and helped analyze the data. BY helped design and perform the SGA experiments. MD and LC helped develop the ABTS overlay assay. CN and CL provided access to critical infrastructure. TM helped design the experiment and analyze the data. The manuscript was written by GS and TM with inputs from RW and additional edits from CN and LC.

## Acknowledgement

The YIp-*TRP1*-natMX plasmid was gifted by Dr. Randy Hampton, the codon optimized *ttLCC1* with a native N-terminal secretion signal was a gift from Dr. Hana Sychrová and the JHY716 query strain was a gift from Dr. Charlie Boone. We thank Ms. Michelle Moksa and Qi Cao from the lab of Dr. Martin Hirst for help with the RT-qPCR experiment, Marjan Barazandeh, Hamid Gaikani and Uche Joseph Ogbede from the Nislow lab for technical help and all members of the Mayor lab for their input and discussion. This work is supported by a NSERC Discovery Grant (RGPIN-2022-03787) and a CFI Innovation Fund (39914).

## Supplementary Material

**Figure S1.**
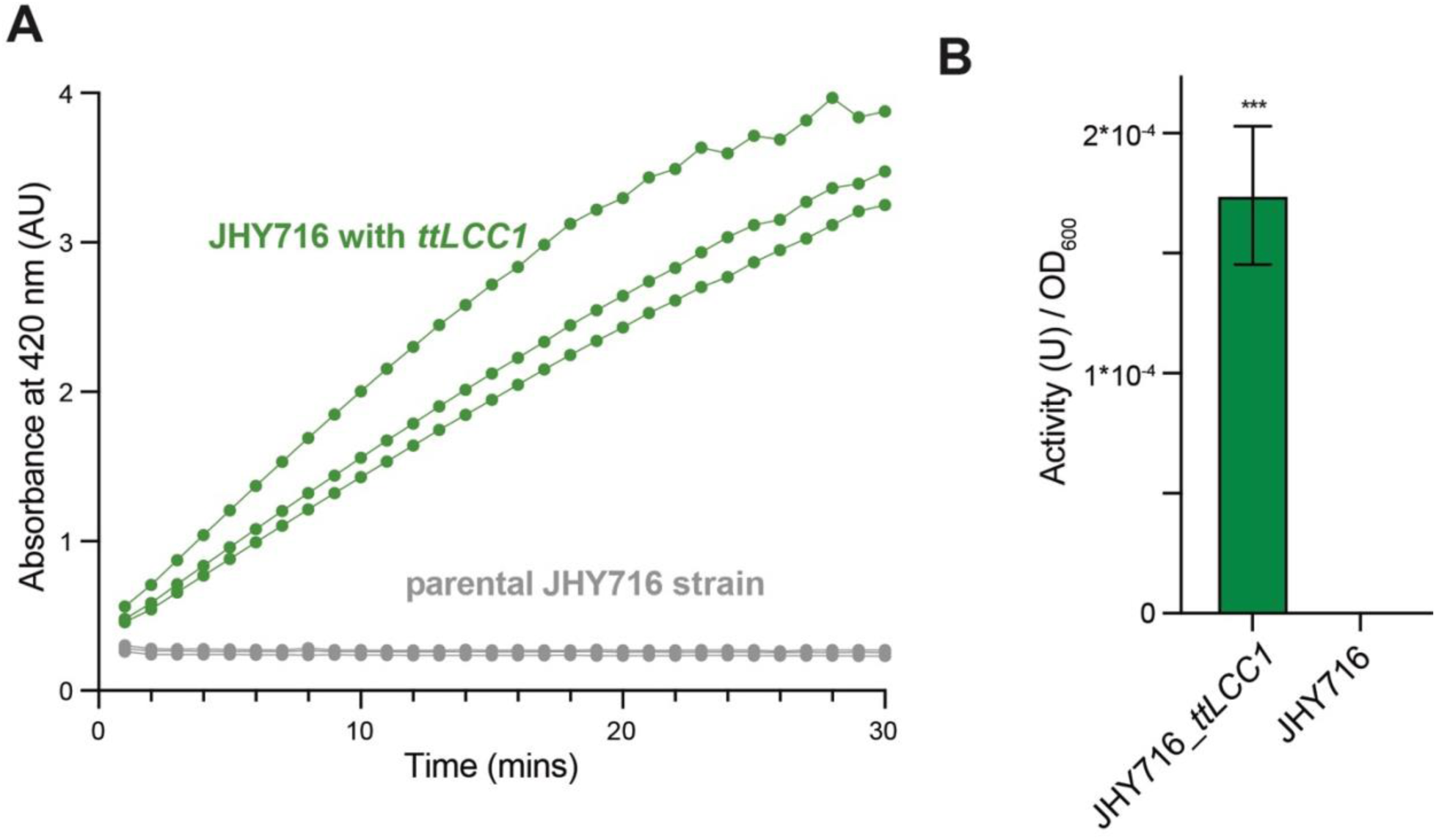
**A and B**. Assessment of secreted laccase activity in liquid cultures. Laccase activity was assessed after collecting the supernatants of the indicated strains following 4 days of growth (three biological replicates). The graph (A) shows the increased signal after the addition of ABTS measured in the plate reader. The bar graph (B) shows the laccase activity (µmols oxidized ABTS / min) normalized to cell density with a p-value of 0.0008 (unpaired t-test).

**Figure S2.**
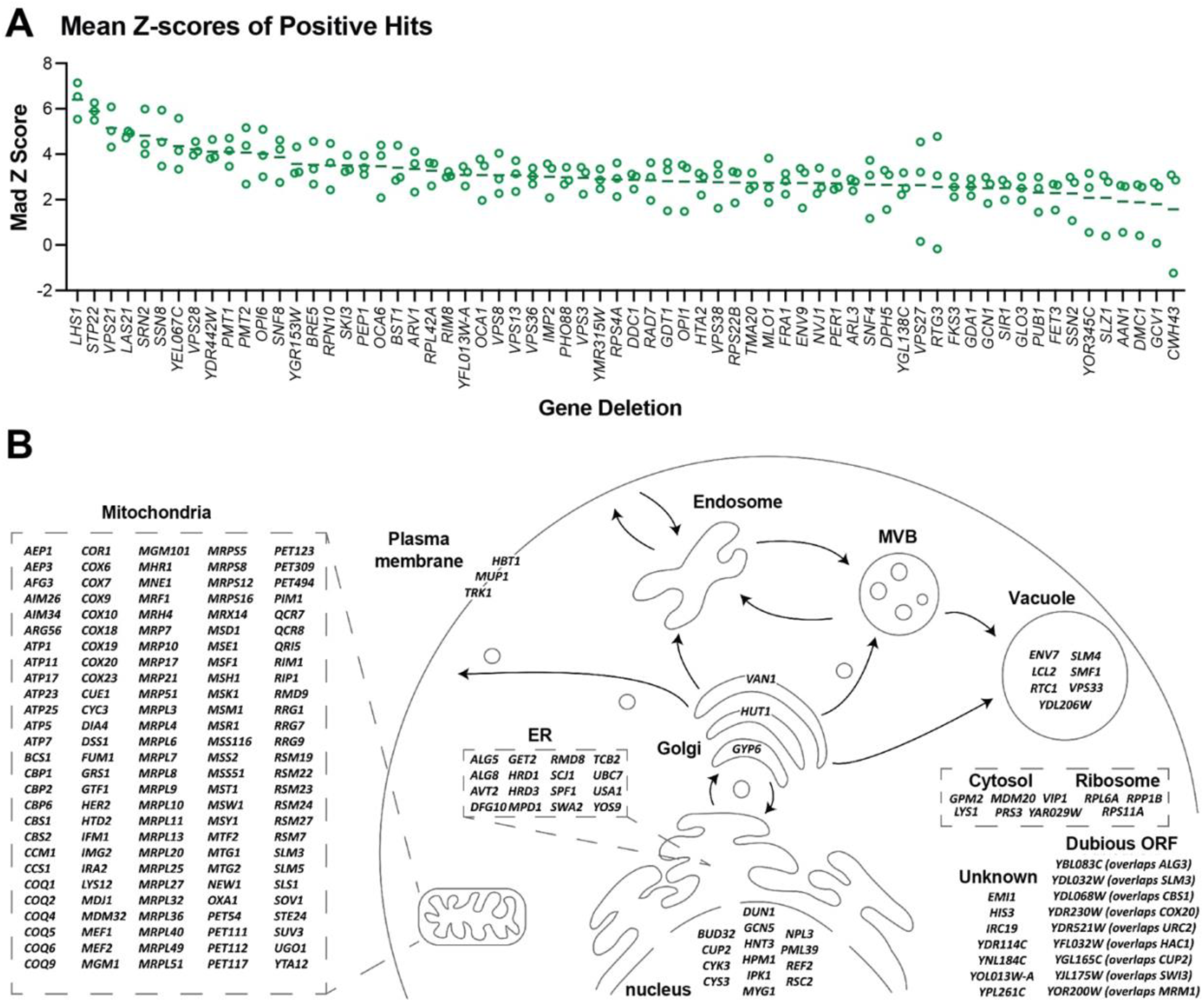
**A**. Ranking of the mean MAD Z-scores of the positive hits from the overlay screen. Circles show the individual MAD Z-scores for each replicate. **B.** Representation of *S. cerevisiae* secretory pathway showing the cellular location of gene deletions that show a decrease in recombinant laccase activity in the ABTS overlay screen.

**Figure S3.**
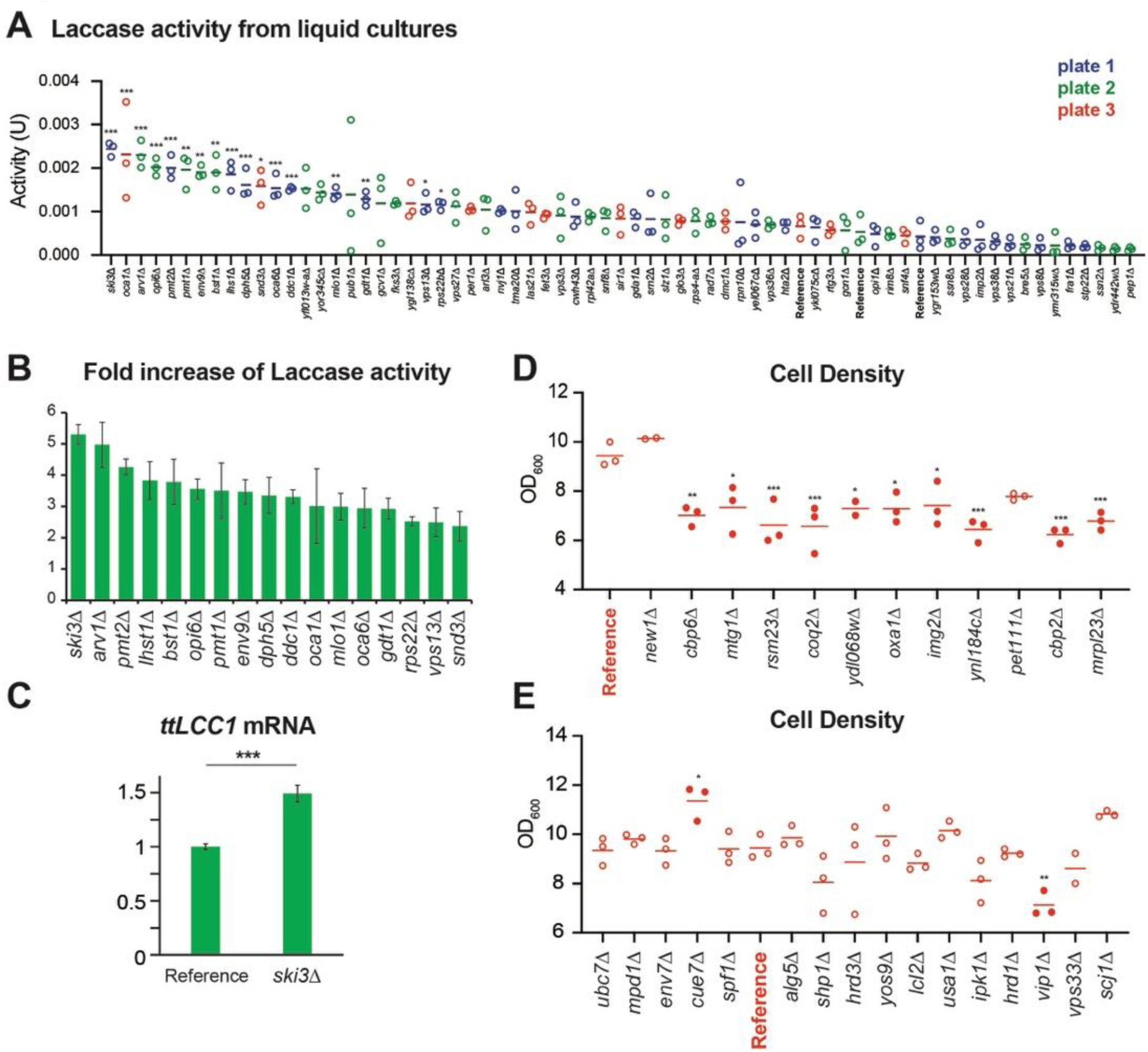
**A.** Laccase activity (µmol/min) of positive hits identified in the overlay screen were assessed in liquid culture. The strains were grown and assessed on three separate sets (plates 1-3). Three biological replicates were grown on the same plate and laccase activity was compared to the parental query strain that was grown on the same plate using ONE way ANOVA tests with Dunnett’s corrections (adjusted p values; < 0.05: *; <0.01: **: <0.005: ***). The normalized activity of these strains is shown in Figure 3A. **B.** Fold change of the normalized laccase activity of the indicated strain in comparison to the reference strain of the cells analyzed in Figure 3A. **C.** Levels of *ttLCC1* mRNA was quantified by RT qPCR in the indicated strains and normalized to *UBC6* mRNA levels using three technical replicates (p-value: 0.004 with a Welch unpaired two tails student t-test). **D and E.** Cell density of a subset of negative hits shown in Figure 3B and C. Three biological replicates for each indicated strain were compared to the reference strain using ONE way ANOVA tests with Dunnett’s multiple comparison corrections.

